# Distinct YFV lineages co-circulated in the Central-Western and Southeastern Brazilian regions from 2015 to 2018

**DOI:** 10.1101/560714

**Authors:** Edson Delatorre, Filipe Vieira Santos de Abreu, Ieda Pereira Ribeiro, Mariela Martínez Gómez, Alexandre Araújo Cunha dos Santos, Anielly Ferreira-de-Brito, Maycon Sebastião Alberto Santos Neves, Iule Bonelly, Rafaella Moraes de Miranda, Nathália Dias Furtado, Lidiane Menezes Souza Raphael, Lucileis de Fátima Fernandes da Silva, Márcia Gonçalves de Castro, Daniel Gaskauskas Ramos, Alessandro Pecêgo Martins Romano, Esper Georges Kallás, Ana Carolina Paulo Vicente, Gonzalo Bello, Ricardo Lourenço-de-Oliveira, Myrna Cristina Bonaldo

**Author notes:** Jointly first authors (Edson Delatorre, Filipe Vieira Santos de Abreu, Ieda Pereira Ribeiro). Jointly last authors (Gonzalo Bello, Ricardo Lourenço-de-Oliveira, Myrna Cristina Bonaldo). **Correspondence to:** Edson Delatorre, /.

## Abstract

The current outbreak of yellow fever virus (YFV) that is afflicting Brazil since the end of 2016 probably originated from a re-introduction of YFV from endemic areas into the non-endemic Southeastern Brazil. However, the lack of genomic sequences from endemic regions hinders the tracking of YFV’s dissemination routes. We assessed the origin and spread of the ongoing YFV Brazilian outbreak analyzing a new set of YFV strains infecting humans, non-human primates (NHP) and mosquitoes sampled across five Brazilian states from endemic and non-endemic regions between 2015 and 2018. We found two YFV sub-clade 1E lineages circulating in NHP from Goiás state (GO), resulting from independent viral introductions into the Araguaia tributary river basin: while the strain from 2017 clustered intermingled with Venezuelan YFV strains from 2000, the YFV strain sampled in 2015 clustered with sequences of the current YFV outbreak in the Brazilian Southeastern region (named YFV_2015-2018_ lineage), displaying the same molecular signature associated to the current YFV outbreak. After its introduction in GO at around mid-2014, the YFV_2015-2018_ lineage followed two paths of dissemination outside GO, originating two major YFV sub-lineages: 1) the YFV_MG/ES/RJ_ sub-lineage spread sequentially from the eastern area of Minas Gerais state to Espírito Santo and then to Rio de Janeiro states, following the Southeast Atlantic basin; 2) the YFV_MG/SP_ sub-lineage spread from the southwestern area of Minas Gerais to the metropolitan region of São Paulo state, following the Paraná basin. These results indicate the ongoing YFV outbreak in Southeastern Brazil originated from a dissemination event from GO almost two years before its recognition at the end of 2016. From GO this lineage was introduced in Minas Gerais state at least two times, originating two sub-lineages that followed different routes towards densely populated areas. The spread of YFV outside endemic regions for at least four years stresses the imperative importance of the continuous monitoring of YFV to aid decision-making for effective control policies aiming the increase of vaccination coverage to avoid the YFV transmission in densely populated urban centers.

## 1 Introduction

In Brazil, the yellow fever virus (YFV) have been sporadically detected in human and non-human primates (NHPs) populations from the enzootic/endemic Northern (Amazon) and epidemic Central-Western regions during the second half of the 20^th^ century (Carrington and Auguste, 2013;Monath and Vasconcelos, 2015). Since the early 2000s, the virus has progressively expanded to the Southeastern and Southern Brazilian regions and in December 2016 began the largest epizootic/epidemic of sylvatic YF registered in the country over the last 50 years (Vasconcelos, 2010;Possas et al., 2018a;Possas et al., 2018b). Between December 2016 and June 2018, a total of 2,139 YF human cases were confirmed in all Southeastern Brazilian states of Minas Gerais (n = 997), São Paulo (n = 577), Rio de Janeiro (n = 307) and Espírito Santo (n = 258) with 735 deaths (case-fatality, 34%). Moreover, in 2019 YFV transmission continues in São Paulo and is emerging in the north of Paraná state from South region of Brazil (Supplementary Figure 1) (Secretaria de Vigilância em Saúde, 2019).

Sequential YF outbreaks reported in the Southeastern and Southern Brazilian regions in 2000– 2001, 2008–2009 and 2016-2018 were more likely caused by single independent events of re-introduction of YFV strains from endemic areas (Mir et al., 2017). A recent study speculated that the YFV strain causing the current outbreak would have been originated in the Brazilian Central-West region. This conjecture was based on the date of the most recent common ancestor of the 2016-2018 Brazilian YFV, which was estimated in a period (July 2014 to January 2016) when YFV circulation was reported in the state of Goiás (Central-Western region) (Rezende et al., 2018). However, the precise routes of dissemination of YFV strains from endemic to non-endemic areas observed in Brazil in the last 15-20 years are difficult to elucidate because of the scarcity of sequences sampled from endemic regions in that period.

The spatiotemporal dynamics of dissemination of the 2016-2018 Brazilian YFV lineage within the Southeastern region also remained unclear. The first study considering full-genome YFV sequences based on samples from Espírito Santo and Rio de Janeiro states from 2017 placed the origin of the Southeastern outbreak in Espírito Santo in April 2016 (July 2015 to October 2016) and supported a rapid southward viral dissemination in direction to the great metropolitan area of Rio de Janeiro (Gomez et al., 2018). A second study also comprising full-genome YFV sequences sampled from Minas Gerais, Espírito Santo and Rio de Janeiro in 2017 supports that the outbreak arose in Minas Gerais in July 2016 (March to November 2016) and was then southerly disseminated toward Espírito Santo and Rio de Janeiro (Faria et al., 2018). A third study, based on partial genome sequences, showed that YFV strains isolated in the state of São Paulo in 2016 branched in basal position relatively to those isolated in Minas Gerais and Espírito Santo at 2017-2018 and traced the origin of current outbreak to July 2015 (July 2014 to January 2016), thus suggesting that the 2016-2018 YFV Brazilian lineage may have circulated in São Paulo state before spread to Minas Gerais and Espírito Santo (Rezende et al., 2018).

Therefore, to define in greater detail the geographic origin and subsequent dissemination routes of the 2016-2018 Brazilian YFV clade, we generated and analyzed 12 YFV new genomes obtained from humans, NHPs and mosquitoes, between 2015 and 2018, from all Southeast Brazilian states (Minas Gerais, Espírito Santo, Rio de Janeiro e São Paulo) and from Central-West region (Goiás). Interestingly, we identified two YFV lineages circulating in Central-West (Goiás) during 2015-2017, one (GO27/2015) displaying the nine unique amino acid signatures characteristic of the 2016-2018 YFV Southeastern outbreak (Bonaldo et al., 2017) and the other (GO05/2017) isolated two year later that displayed an amino acid pattern typical of older YFV sub-clade 1E strains sampled in Brazil and Venezuela between 2000 and 2010 (de Souza et al., 2010). Besides, these 12 new YFV complete coding region sequences (CDS) were combined with previously described YFV CDS from Brazil (n = 67), Venezuela (n = 5) and Trinidad and Tobago (n = 1) and then subjected to phylogeographic analyses to get a better picture of the ongoing YFV epidemic in Brazil.

## 2 Materials and Methods

### 2.1 YFV samples

Viral samples from 12 infected hosts from distinct biomes and river basins in the states of Goiás (*n* = 2), Central-Western region, and Rio de Janeiro (*n* = 2), Minas Gerais (*n* = 3), Espírito Santo (*n* = 3) and São Paulo (*n* = 2), Southeastern region of Brazil, were analyzed (Table 1). Serum samples of human cases, liver samples of NHPs and homogenates of entire bodies of pooled adult female mosquitoes were collected and processed as previously described (Ferreira-de-Brito et al., 2016;Bonaldo et al., 2017;Gomez et al., 2018;Abreu et al., 2019). The YFV isolates ABR1005 and ABR1009 were obtained from the sera of a 64 and 30-year-old male patients, respectively. The ABR1005 patient was hospitalized four days after onset of symptoms and died due to multiple organ failure. The ABR1009 patient was hospitalized four days after onset of symptoms, but fully recovered from the infection. These two subjects were included in a study protocol approved by the institutional review boards at the Hospital das Clínicas (School of Medicine, University of São Paulo) and the Infectiology Institute Emilio Ribas (CAAE: 59542216.3.1001.0068). The analysis of human samples was carried out in accordance with the recommendations of the Ethics Committees for human research at Instituto Oswaldo Cruz (CAAE 69206217.8.0000.5248), which exempted the need of a specific written informed consent from patients or their legal representatives. Capture of NHPs and mosquitoes, as well as management of NHP samples, were carried out in accordance with the Brazilian environmental authorities (SISBIO-MMA licenses 54707-6 and 52472-2, and INEA licenses 012/2016, 019/2018) and Ethics Committee of Animal use at Instituto Oswaldo Cruz (IOC) (CEUA license L037/2016).

**Table 1:**
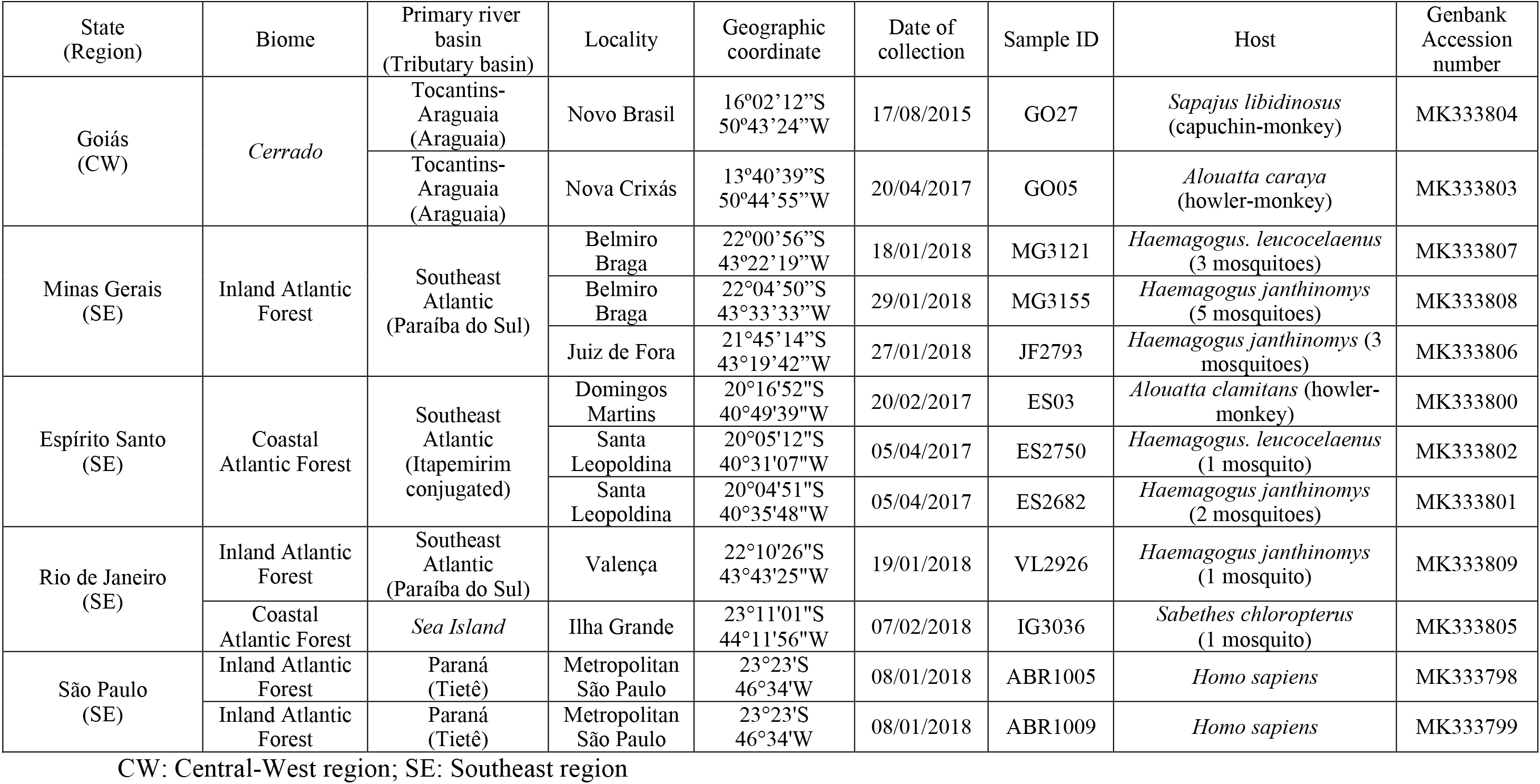
YFV samples from Brazil sampled in 2015, 2017 and 2018.

### 2.2 YFV genome detection and Nucleotide Sequencing

The sera from the individuals from São Paulo state (ABR1005 and ABR1009) and an NHP liver homogenate sample from Goiás (GO27) were employed to obtain first-passage YFV isolates by infection of monolayer cell cultures of the C6/36 clone of *Aedes albopictus*. Viral RNA was obtained from cell cultures or directly from samples as described elsewhere (Bonaldo et al., 2017). The set of primers utilized in PCR and sequencing procedures followed a previous report (Gomez et al., 2018). Nucleotide sequences were determined by capillary electrophoresis at the sequencing facility of Fiocruz-RJ (RPT01A – Sequenciamento de DNA – RJ). The sequences were assembled with SeqMan Pro version 8.1.5 (DNASTAR, Madison, WI, USA). The Molecular Evolutionary Genetics Analysis (MEGA) 7.0 program (Kumar et al., 2016) was adopted to explore the amino acid differences as well as to calculate nucleotide and amino acid distances.

### 2.3 Evolutionary and Phylogeographic analyses

Complete CDS (10,239 nt in length) of the 12 newly generated YFV genomes were combined with CDS of YFV American sequences available in GenBank (www.ncbi.nlm.nih.gov) according to the following inclusion criteria: 1) link to a publication, 2) coverage of at least 99% of the viral CDS, and 3) known date and country of collection. The sequences were aligned with MAFFT (Katoh and Standley, 2013) and viral phylogenies were reconstructed by maximum likelihood (ML) analysis implemented in PhyML (Guindon et al., 2010) applying the best substitution model selected by jModelTest v1.6 (Darriba et al., 2012). The temporal signal of different combinations of sequences representing the South American genotypes I and II (SA-I+II), the South American genotype I (SA-I), and the Modern lineage of SA-I (Mir et al., 2017) were examined with TempEst v1.5.1 (Rambaut et al., 2016). The dataset with the best temporal structure (SA-I) was chosen for the subsequent time tree reconstructions by Bayesian method (Supplementary Table 1).

The time scale of the SA-I and 2016-2018 Brazilian YFV datasets were estimated using the Markov chain Monte Carlo (MCMC) algorithms implemented in the BEAST v1.8.4 package (Drummond and Rambaut, 2007;Drummond et al., 2012) with BEAGLE (Ayres et al., 2012) to improve running time. The evolutionary process was estimated using the best-fit nucleotide substitution model (GTR+I+Γ4), a relaxed uncorrelated lognormal molecular clock model (Drummond et al., 2006) and the non-parametric Bayesian Skyline coalescent tree prior (Drummond et al., 2005). A CTMC rate reference prior (Ferreira and Suchard, 2008) and a normal prior (mean = 4.5×10^−4^ substitution/site/year, standard deviation = 1.0×10^−4^) in the evolutionary rate were used for the analysis of the SA-I and 2016-2018 Brazilian YFV datasets, respectively.

The reconstruction of migration events throughout the phylogeny for the 2016-2018 Brazilian YFV lineage also employed the BEAST package using discrete and continuous models. The discrete phylogeographic analysis was performed using reversible (symmetric) and nonreversible (asymmetric) discrete phylogeographic models (Lemey et al., 2009), assigning discrete traits for each sequence representing the Brazilian state of isolation, except for MG, in which further geographic subdivision were employed. The spatiotemporal reconstruction in continuous space utilizing the geographic coordinates (latitude and longitude) of each YFV isolate was estimated with a homogenous Brownian diffusion (BD) model and the heterogeneous Cauchy, Gamma and Lognormal relaxed random walk (RRW) models (Lemey et al., 2010). Comparisons among the different discrete and continuous phylogeographic models were performed using the log marginal likelihood estimation (MLE) based on path sampling (PS) and stepping-stone sampling (SS) methods (Baele et al., 2012). Bayesian analyses were run for 10^8^ generations and convergence (effective sample size > 200) was inspected using TRACER v1.7 (Rambaut et al., 2018) after discarding 10% burn-in. The maximum clade credibility (MCC) trees were summarized using TreeAnnotator v.1.8.4 (Drummond et al., 2012) and visualized with FigTree v.1.4.4 (http://tree.bio.ed.ac.uk). The viral spatio-temporal diffusion was analyzed and visualized in SPREAD (Bielejec et al., 2011) and further projected in maps generated with QGIS software (http://qgis.org) using public access data collected from the Brazilian Institute of Geography and Statistics (IBGE, 2019) and National Water Agency (ANA, 2019).

## 3 Results

### 3.1 Identification of two YFV lineages circulating in the state of Goiás during 2015-2017

Initially, we sequenced the complete genomes of YFV obtained at two Brazilian biomes to infer the origin and dissemination routes of YFV during the 2016-2018 outbreak in the country. Ten YFV strains were sampled from human, NHP and mosquitoes in all Southeastern Brazilian states: Minas Gerais, Espírito Santo, Rio de Janeiro and São Paulo, located in the Atlantic Forest biome and where massive epizootics and human cases occurred. Two other YFV genomes were recovered from NHP sampled in Goiás, a state of the Brazilian Central-Western region whose predominant vegetation is the *Cerrado*, a savanna-like biome occupying the territory between the Amazon and Atlantic rain forest of the Southeast region (Table 1). Both YFV strains from Goiás were sampled from sites located in the Tocantins-Araguaia river basin. The ML and Bayesian phylogenetic analyses placed the newly generated YFV genomes inside the sub-clade 1E (de Souza et al., 2010) of the Modern lineage of SA-I (Mir et al., 2017), with high support [aLRT/posterior probability (*PP*) = 1] (Figure 1 and Supplementary Figure 2). One virus sample from Goiás (GO27/2015) infecting a capuchin-monkey from Novo Brasil on August 2015, clustered in a highly supported (*PP* = 1) clade (YFV_2015-2018_) with all Brazilian YFV sequences from the Southeastern region associated with the current outbreak. By contrast, the other strain from Goiás (GO05/2017) that infected a howler-monkey from Nova Crixás on April 2017, was intermingled among Venezuelan YFV genomic sequences from the 2000s. The mean evolutionary rate for the YFV SA-I was estimated at 4.6×10^−4^ substitution/site/year (Supplementary Table 2), fully consistent with that previously reported (Nunes et al., 2012;Gomez et al., 2018), while the time of the most recent common ancestor (T_MRCA_) of the YFV_2015-2018_ clade was estimated on December 2013 [95% Bayesian credible interval (BCI): August 2012 to December 2014].

**Figure 1.**
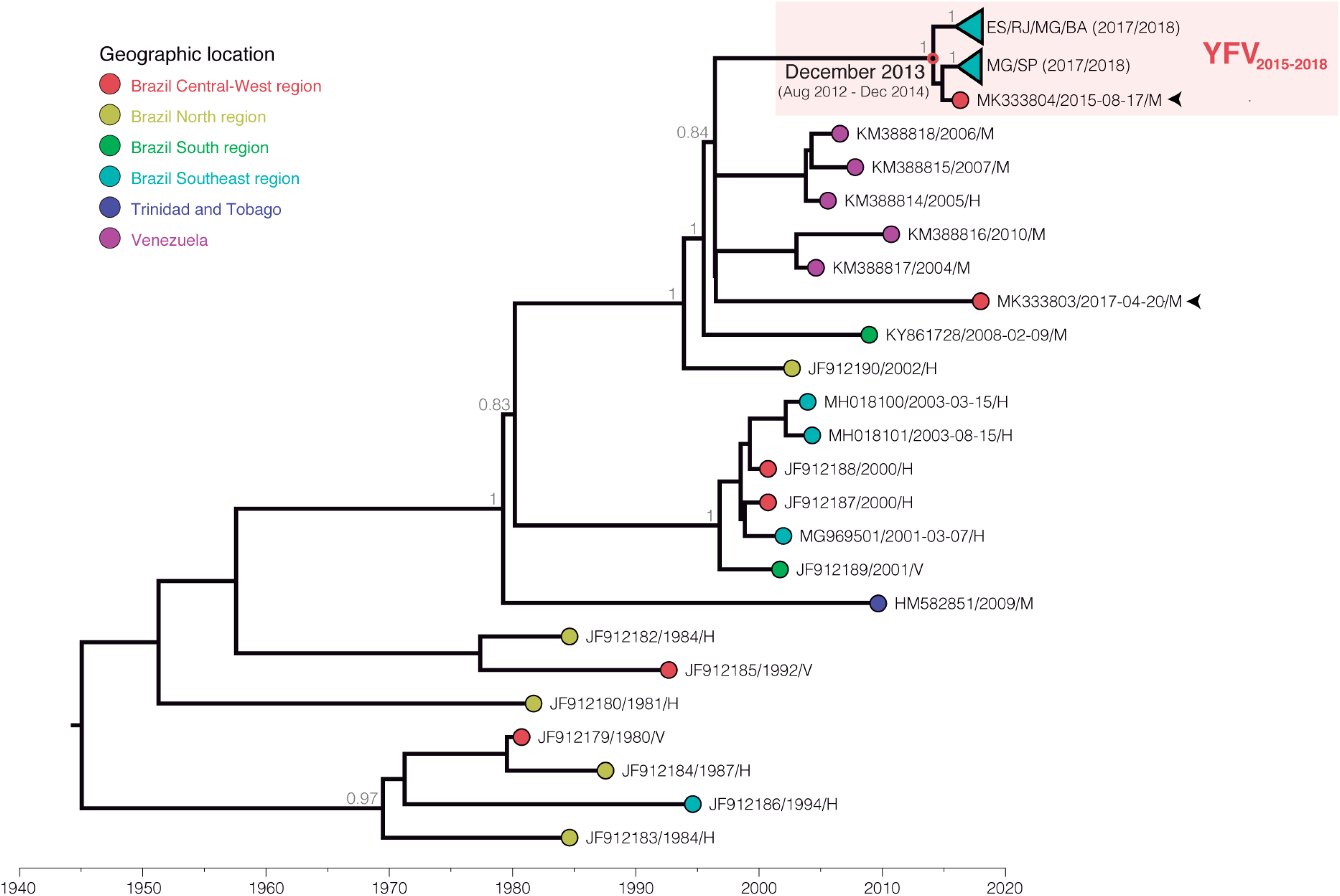
Bayesian maximum clade credibility phylogeny of the YFV South American genotype I. Only posterior probabilities (*PP*) of key nodes are denoted. The external node color indicates sample location (geographic region or country) accordingly to the legend. The samples obtained from NHP from Goiás state are indicated by arrows. The clade comprising the GO217/2015 sample and all YFV genomes from the ongoing outbreak (YFV_2015-2018_) is highlighted by a red box, and inner clades were collapsed for clarity. The node representing the most recent common ancestor of YFV_2015-2018_ lineage is indicated by a red circle along with its temporal origin estimate. All horizontal branch lengths are drawn to a scale of years. Sequences names are coded as accession number/date of collection/host (H– human, V – mosquito vector, M – NHP.

Analysis of amino acid signatures showed the ubiquitous presence of the nine unique amino acid substitutions previously described (Bonaldo et al., 2017;Gomez et al., 2018) in almost all new YFV Brazilian genomes from the Southeastern region and in one sample from Goiás (GO27/2015) (Supplementary Table 3 and Supplementary Figure 3). Distinctly, the other YFV strain from Goiás (GO05/2017), exhibited an amino acid pattern more similar to older YFV sub-clade 1E strains sampled in Brazil and Venezuela between 2000 and 2010 (Supplementary Table 3 and Supplementary Figure 3). These results clearly support the circulation of at least two YFV sub-clade 1E lineages in the *Cerrado* between 2015 and 2017 that probably resulted from independent viral introductions in the Araguaia tributary basin of the Tocantins-Araguaia primary watershed. Moreover, these results revealed that the molecular signature previously associated to the 2016-2017 YFV Southeastern Brazilian strains was already present in a YFV strain isolated in the *Cerrado* biome in 2015.

### 3.2 The YFV_2015-2018_ lineage likely arose in Goiás in 2014 and was disseminated to the Southeastern region following two major routes

To determine with more precision the geographic origin and dissemination routes of the YFV_2015-2018_ lineage, we first applied discrete Bayesian symmetric and asymmetric phylogeographic models. The Bayes Factor test showed no significant support in favor of one of the discrete phylogeographic models (Supplementary Table 4), thus we presented the resulting phylogeographic reconstruction of the simpler symmetric model (Figure 2). According to this analysis, the YFV_2015-2018_ lineage likely originated in the state of Goiás [posterior state probability (*PSP*) = 0.44] in June 2014 (95% BCI: January 2013 to June 2015). From there, it followed two paths of dissemination towards the state of Minas Gerais, originating two major YFV sub-lineages in the Southeastern region here called YFV_MG/ES/RJ_ and YFV_MG/SP_.

**Figure 2.**
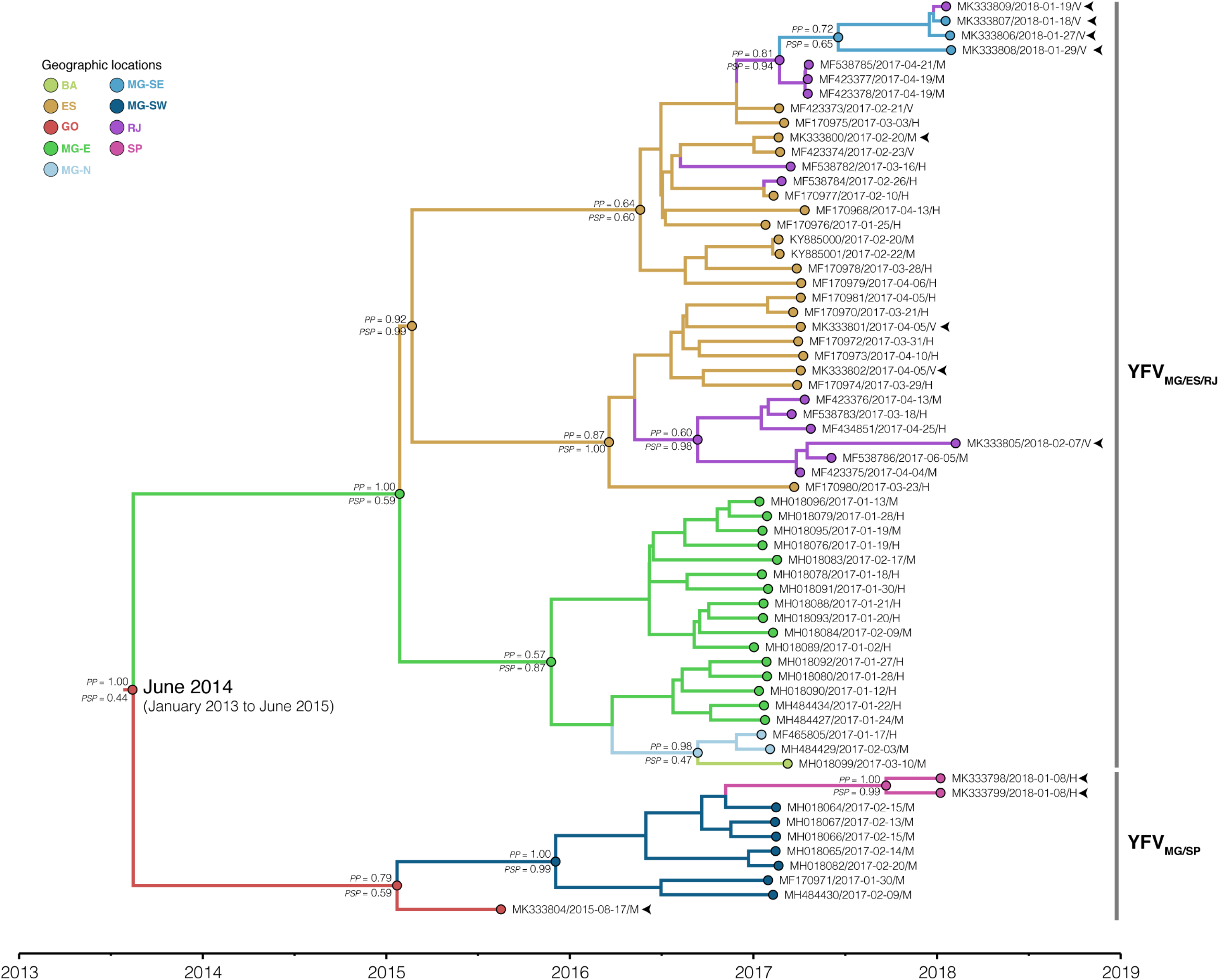
Time-scaled maximum clade credibility phylogeny of the YFV_2015-2018_ lineage. The branches’ colors represent the most probable location of their descendent nodes as indicated in the legend. The posterior (*PP*) and posterior state (*PSP*) probabilities are denoted above and below key branches, respectively. The time for the most common ancestor of lineage YFV_2015-2018_ is indicated at the basal node (red circle) with the 95% HPD in parenthesis. Tips names were codified as accession number/date/region/host. The tips corresponding to the samples sequenced in this study are indicated by arrows. All horizontal branch lengths are drawn to a scale of years. BA, Bahia; ES, Espírito Santo; GO, Goiás; MG-E, Eastern Minas Gerais; MG-N, Northern Minas Gerais; MG-SE, Southeastern Minas Gerais; MG-SW, Southwest Minas Gerais; RJ, Rio de Janeiro; SP, São Paulo.

The YFV_MG/ES/RJ_ sub-lineage probably reached initially the eastern region of Minas Gerais (MG-E, *PSP* = 0.59) on December 2015 (95% BCI: May 2015 to June 2016), from where it most likely spread to Espírito Santo state (*PSP* = 0.99) on April 2016 (95% BCI: December 2015 to September 2016) and to the northern area of Minas Gerais (MG-N, *PSP* = 0.47) on June 2016 (95% BCI: January 2016 to October 2016). From MG-N, the YFV spread to the south of the state of Bahia, in the Northeastern Brazilian region. From ES, the YFV was introduced in Rio de Janeiro at least four times, generating two successful intrastate transmission chains that advanced southwards: 1) one chain followed the northern side of the Serra do Mar along the Paraíba do Sul tributary basin, surpassing the Rio de Janeiro metropolitan region towards the south (Valença municipality) and affecting the southeastern region of Minas Gerais (MG-SE, *PSP* = 0.65); 2) the other transmission chain spread through the coastal area South of Serra do Mar, crossing the Rio de Janeiro state metropolitan region and reaching an island located in the southern region of Rio de Janeiro (Ilha Grande).

The YFV_MG/SP_ sub-lineage most likely spread from Goiás into the southwestern area of Minas Gerais (MG-SW, *PSP* = 0.99) on April 2016 (95% BCI: October 2015 to September 2016) and then disseminated from MG-SW to the metropolitan region of the state of São Paulo. Remarkably, four YFV genomes from MG-E and one from MG-SW displayed differences in the molecular amino acid signature (D15762E; R2607Q). Nevertheless, the variations were not fixed in 2018 YFV samples as identified in the position I2176V clustering in YFV_MG/ES/RJ_, which were collected in Vale do Paraíba basin in 2017 and 2018, suggesting the maintenance of the YFV polymorphism in this region. Overall, these results expanded the geographical and temporal edge of the YFV_2015-2018_ lineage and further revealed a very low degree of phylogenetic intermixing of YFV_2015-2018_ strains from different Brazilian states during viral dissemination in the Southeastern region.

### 3.3 The YFV_2015-2018_ lineage was disseminated following primary river basins in the Southeastern region

To get some insight regarding the spatiotemporal dynamics of dissemination of the YFV_2015-2018_ lineage, we applied different continuous phylogeographical models, assuming homogeneous (BD) and heterogeneous (RRW) dispersion rates among lineages. The RRW model with Gamma distribution was strongly supported as the fittest diffusion model (Supplementary Table 5), indicating significant variation in the diffusion rate among the branches. The phylogeographic continuous model (Figure 3) changed slightly the epicenter of the YFV_2015-2018_ lineage but supports the existence of two main routes of dissemination within the Brazilian Southeastern region and few viral migrations between different states, consistent with the discrete phylogeographic reconstruction. This analysis also supports that the YFV was disseminated following major river basins.

**Figure 3.**
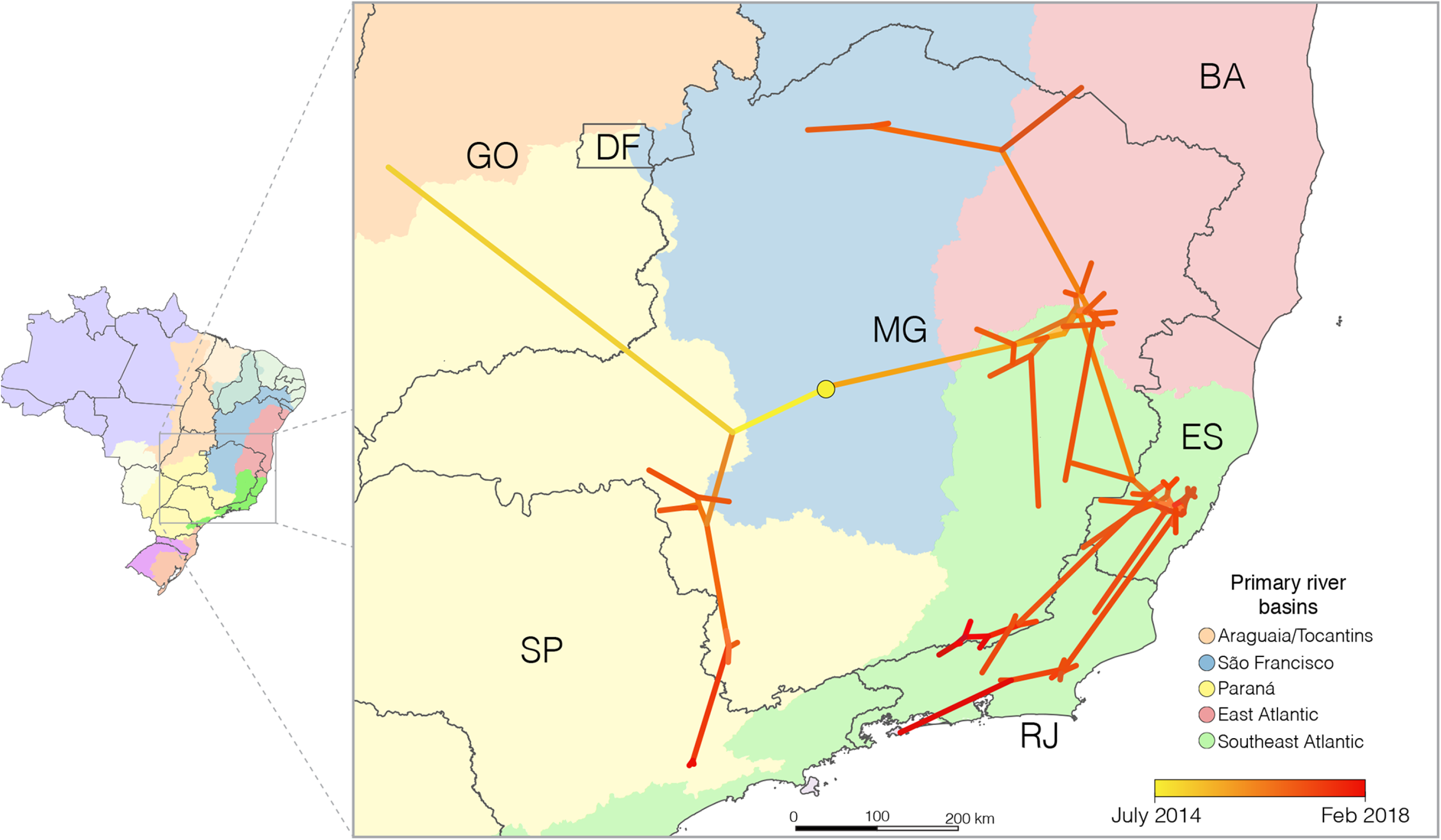
Reconstructed spatiotemporal diffusion of the YFV_2015-2018_ lineage. Phylogeny branches were arranged in space according the internal nodes locations inferred by the continuous phylogeographic model. Branches were colored according to time as indicated by the legend. The dark gray lines represent the Brazilian states boundaries while the colored areas the different primary river basins, as indicated. Brazilian states: DF, Distrito Federal; ES, Espírito Santo; GO, Goiás; MG, Minas Gerais; RJ, Rio de Janeiro; SP, São Paulo.

According to the continuous model, the origin of the YFV_2015-2018_ lineage would be the central region of Minas Gerais, within the São Francisco watershed, from where it would have independently spread to the west, east and south, reaching the Tocantins-Araguaia basin in the Goiás in 2015, the Southeast Atlantic hydrographic region in Minas Gerais at the beginning of 2016, and the Paraná hydrographic region of that state in the middle of 2016, respectively. From the eastern of Minas Gerais, the virus moved southward following the Southeast Atlantic watersheds distributed among the states of Minas Gerais, Espírito Santo, and Rio de Janeiro (an area covered by the Atlantic forest biome) and northward, returning to the São Francisco river basin and south of Bahia. Simultaneously, the YFV lineage showed southerly dissemination from the southwestern region of Minas Gerais toward the São Paulo state, following the Paraná basin. We estimate that YFV lineages moved, on average, 0.8 km/day (95% BCI: 0.5 to 1.0 km/day).

## 4 Discussion

The current re-emergence of YFV in the Southeastern Brazilian region resulted in the largest outbreak of sylvatic YF observed in South America in the last decades. The transmission has been expanding southward in Brazil reaching sites considered YFV-free areas for 80 years, and therefore with scanty YFV vaccination coverage. As a result, 38 cases and nine deaths have been reported in January 2019, around 1,160 km from the first signal of increased incidence of YF in the Southeast (north-western region of Minas Gerais) in late 2016 (Secretaria de Vigilância em Saúde, 2019).

Previous studies pointed out that the ongoing YFV outbreak in the Brazilian Southeastern region resulted from a single introduction event of a YFV Modern lineage strain from an endemic area (Mir et al., 2017;Faria et al., 2018;Gomez et al., 2018;Rezende et al., 2018). However, the precise route of viral dissemination was not achieved due to the scarcity of Brazilian YFV sequences sampled from endemic regions over the last years. Here, the analysis of an NHP YFV sample from Goiás (Central-Western region) in 2015 revealed that it is phylogenetically related with and carry the same amino acid signature of the YFV strains causing the current outbreak in the Southeastern region. Moreover, the discrete phylogeographic analysis showed that the YFV causing the current Brazilian outbreak probably originated in Goiás at around mid-2014, a result congruent with epidemiological reports on human NHP infections (Supplementary Figure 1). Altogether, our data stressed the origin of the current YFV outbreak in the Central-western region and expanded the estimated T_MRCA_ of the ongoing YFV outbreak to almost two years before it gained epidemiological visibility in the end of 2016 (Secretaria de Vigilância em Saúde, 2017). Interestingly, we also identified for the first time, two YFV lineages circulating in Goiás during 2015-2017, one of which followed two independent paths of dissemination towards Minas Gerais, originating two major YFV sub-lineages in the Southeast region responsible for the severe ongoing outbreak.

The discrete phylogeographic analysis pointed out that the current YFV Brazilian outbreak probably originated in Goiás, while the continuous phylogeographic model placed the epicenter in the central region of Minas Gerais state. The paucity of YFV sequences from the *Cerrado* biome of the state of Goiás probably explained such discordant results. The placement of the root location of the YFV_2015-2018_ lineage in Goiás is more consistent with the official reports of the Brazilian Ministry of Health (Secretaria de Vigilância em Saúde, 2015). According to the spatiotemporal epidemiologic reports of YFV infections in both humans and NHPs in the national surveillance system, after five years of no records of YFV cases outside the Amazon, 31 NHPs died between May and August 2014 in seven municipalities in the Tocantins state, in the Tocantins-Araguaia basin. The only corpse found still adequate to diagnosis was positive to YFV (Secretaria de Vigilância em Saúde 2014). The Tocantins-Araguaia basin drains the territory of Tocantins and the great northern part of the neighbor state of Goiás into de Amazon river, and the gallery forests along its tributaries consist of a large network of corridors between the Amazon and *Cerrado* biomes. Thus, soon in early 2015, besides in Tocantins, YFV infections were detected in Goiás, and the virus spilled over from the Tocantins-Araguaia into the São Francisco and Paraná basins, with reports scattered in northwestern Minas Gerais and southern Goiás and the Federal District (Brasília) (Figure 4).

**Figure 4.**
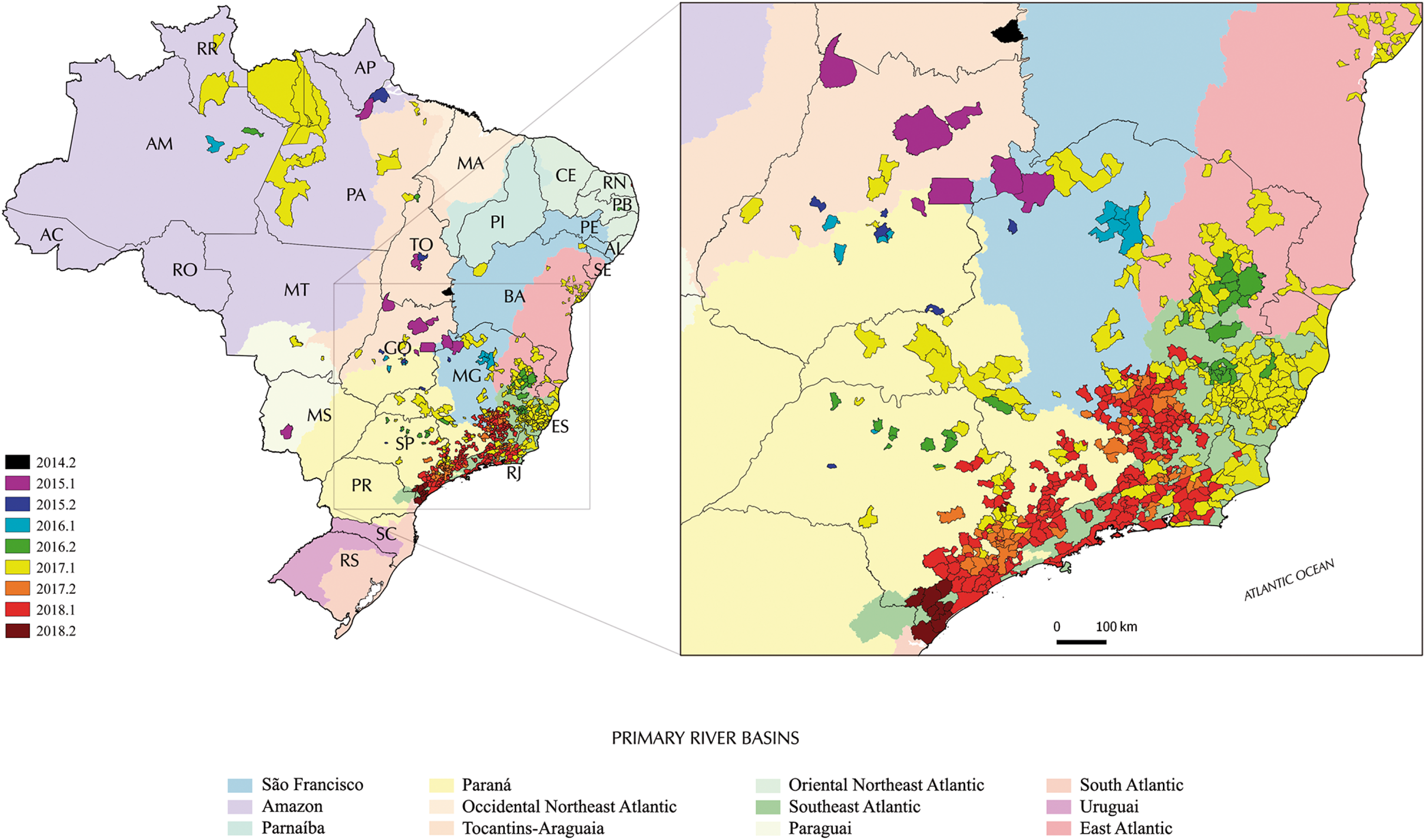
Municipalities reporting YFV in Brazil from late 2014 to late 2018 according to epidemiological data. Colors distinguish the semester when the first confirmed human and/or non-human YFV infection was recorded in such a municipality in a half-yearly basis during the period. Brazilian states: AC, Acre; AL, Alagoas; AP, Amapá; AM, Amazonas; BA, Bahia; CE, Ceará; DF, Distrito Federal; ES, Espírito Santo; GO, Goiás; MA, Maranhão; MT, Mato Grosso; MS, Mato Grosso do Sul; MG, Minas Gerais; PA, Pará; PB, Paraíba; PR, Paraná; PE, Pernambuco; PI, Piauí; RJ, Rio de Janeiro; RN, Rio Grande do Norte; RS, Rio Grande do Sul; RO, Rondônia; RR, Roraima; SC, Santa Catarina; SP, São Paulo; SE, Sergipe; TO, Tocantins.

Brazilian Health authorities consider this YFV reemergence as the start of the north-southeast virus spreading wave that has not stopped yet (Secretaria de Vigilância em Saúde, 2015;2017;2019). Accordingly, in late 2015 and early 2016, the circulation of YFV was detected in sites of Minas Gerais and Goiás, but still essentially limited to the *Cerrado* biome in the three above mentioned river basins. Intriguingly, in late 2016, human cases and epizooties due to YFV were recorded simultaneously and independently in the northeast Minas Gerais as well as in southeastern Minas Gerais/northern São Paulo, where the virus has then gained the Atlantic forest ecosystem. In fact, the outbreak was officially recognized only when the epidemiological records rapidly peaked and as that the virus reached the Southeast Atlantic river basin in Espírito Santo and Rio de Janeiro, as well as continued to spread in all primary watershed of São Paulo and Minas Gerais throughout 2017. One of the causes of this delay was the introduction of Chikungunya (2014) and Zika (2015) virus in Brazil and the resulting epidemics, which probably reduced sensitivity of surveillance and interfered with the visibility of the YFV reemergene. In 2017, scattered records of YFV circulation were also made in the states of Bahia, Goiás, Mato Grosso and even in the Amazon. In early 2018, epidemiological data suggested that transmission was mostly concentrated in the Southeast Atlantic basin in the states of Rio de Janeiro, Minas Gerais and São Paulo and eastern Paraná basin (Figure 4). Then, the place and time of origin for the current YFV Brazilian outbreak here estimated fully agree with the above mentioned officially confirmed infections in NHP and humans reported by the Brazilian Ministry of Health (Secretaria de Vigilância em Saúde, 2015). Indeed, the Novo Brasil municipality in Goiás, from where the NHP YFV sample analyzed here was taken, is located about 100 km from the banks of the Araguaia river. The earlier detection of YFV in NHP from the Tocantins indicates that the state of Goiás probably acted as a staging post during the dissemination of the YFV_2015-2018_ lineage from the North (Amazon basin) to the Southeast Brazilian regions. In this sense, the Tocantins-Araguaia watersheds may have played a major role in YFV dissemination as it extends from more than 2.500 km following two river axes (Araguaia and the Tocantins), offering a contiguous connection between the *Cerrado* (southward, Goiás and Tocantins states) and Amazon (northward, Pará state) biomes (ANA, 2019).

Both discrete and continuous phylogeographic models combined with the epidemiological records support of the YFV_2015-2018_ lineage moved toward densely populated Southeastern urban regions with low YFV vaccine coverages following major routes along different primary river basins. The phylogeographic analyses pointed out that the YFV_2015-2018_ lineage probably arrived in the Southeast region via the São Francisco watershed located in Minas Gerais and then moved to the Southeast Atlantic watersheds in the east and the Paraná hydrographic region in the southwest. The viral lineage that moved following the Southeast Atlantic watersheds reached the eastern and northern areas of Minas Gerais state, as well as the south of Bahia, Espírito Santo and Rio de Janeiro states. The viral lineage that followed the Paraná hydrographic region spread to the Southwest of Minas Gerais and São Paulo states. A previous study proposed that the YFV_2015-2018_ lineage was introduced in the southeastern region through São Paulo and then moved to other Southeastern states (Rezende et al., 2018). However, this conclusion is hampered as they used only partial genomes (with a very low number of nucleotide substitutions supporting the phylogenetic relationships) and did not conduct a formal phylogeographic analysis. Although Faria et al. (2018) already described that the YFV_2015-2018_ lineage displayed southward and eastward expansion from its inferred origin in Minas Gerais, our results consist of the first description of concurrent dispersion of the YFV_2015-2018_ lineage following two independent routes that seems to be linked to the main hydrographical basins.

Our phylogeographic analyses also support that the rapid spread of the YFV_2015-2018_ lineage in the Southeastern region seems to have resulted from a few successful viral disseminations events between states. Most YFV transmission in Espírito Santo was probably originated from a single successfully transmission from Minas Gerais, while most viral transmissions in Rio de Janeiro seems to have resulted from two independent introductions from Espírito Santo that subsequently spread along the coastal and northern sides of the Serra do Mar mountain system, as previously described (Gomez et al., 2018). We found that both transmission chains previously detected in Rio de Janeiro state continued to expand to beyond the metropolitan region, reaching municipalities close to the border with the state of São Paulo during 2018. The two YFV 2018 genomes from São Paulo analyzed here are the first described of the current YFV outbreak from that state and were the result of independent dissemination from southwestern region of Minas Gerais, but more sequences from São Paulo are necessary to understand the epidemic dynamic in this state. It is unclear if most YFV infections in São Paulo resulted from a single or a few founder viral strains that spread from the southwest of Minas Gerais along the Paraná hydrographic basin, or if other viral strains may have also been disseminated from Rio de Janeiro along the Southeast Atlantic watersheds. The recent detection of YFV in NHP from the coastal area of Paraná state in 2019 (Secretaria de Vigilância em Saúde, 2019) indicates continuous dissemination of YFV into the Southern region probably following the Paraná and/or the Southeast Atlantic hydrographic basins.

We estimated that the YFV_2015-2018_ lineage moved on average 0.8 km/day and similar results were obtained when the YFV_2015-2018_’s outgroup sequence GO27/2015 was removed from the analysis. This velocity is lower than the estimates described by Faria et al. (2018) and Gomez et al. (2018) that also analyzed the dispersion of the ongoing YFV outbreak in Brazil and found dispersion rates of 4.2 and 3.4 km/day, respectively. However, those studies analyzed sequences sampled between January-April of 2017, corresponding to the wet and warmer season, when there is an increase in the density of vectors (Alencar et al., 2018) facilitating the transmission. The primary vectors in the current outbreak in Southeast Brazil are the mosquitoes *Haemagogus leucocelaenus* and *Haemagogus janthinomys* (Abreu et al. 2019), which can disperse large distances in short time (Causey et al., 1950). Our conservative velocity of YFV dispersion fully agrees with the observed distances traveled by mosquitoes, but also with howler monkeys in Southeastern Brazil (Jung et al., 2015) and also agrees with an epidemiological model based on dates and place of reported monkey deaths, which estimated YFV displacement speeds of 2.7 km/day in the warmer months and 0.5 km/day in the coldest months (Fioravanti, 2018). Thus, the YFV dispersion velocity estimated in this study would correspond to a median value between these two speeds.

Surprisingly, we found two YFV sub-clade 1E lineages circulating in the Araguaia tributary basin, indicating at least two independent introductions of YFV in that region probably from the enzootic/endemic Amazon biome in a narrow time frame. While sample GO27/2015 isolated in 2015 displays the nine unique amino acid signatures characteristic of the 2016-2018 YFV Southeastern outbreak, sample GO05/2017 isolated two years later at the same watershed exhibited an amino acid pattern typical of older YFV sub-clade 1E strains sampled in Brazil and Venezuela between 2000 and 2010. According to our analysis, only YFV strains related to the GO27/2015 were able to further disseminate from Goiás towards the Southeast region. We can speculate that the different pattern of molecular signatures present in the two YFV strains from Goiás modulate the spread of each viral lineage since some of them were located in key viral enzymes (Gomez et al., 2018). Alternatively, the GO05/2017 lineage may have been introduced from the Amazon into Goiás at a later time, and its dissemination towards the Southeast was hampered due to the reduction or even exhaustion of susceptible NHP hosts caused by the previous passage of the lineage that originated the YFV_2015-2018_ clade. Consistent with this last hypothesis, a recent ecological study concludes that dissemination of YFV in South America is not random, but it is influenced by key geo-environmental factors like diversity and number of susceptible NHP hosts (Hamrick et al., 2017). Curiously, epidemiological records showed YFV transmission in several sites in the Amazon in early 2017, including in the Tocantins-Araguaia basin in Pará state (Figure 4)

In summary, we showed that at least two different YFV lineages circulated in the *Cerrado* biome (Araguaia tributary basin) in a narrow time frame. One of these lineages further spread out the *Cerrado* biome in Goiás to the Atlantic forest biome in the Brazilian Southeastern region, originating the current Brazilian outbreak (YFV_2015-2018_ lineage) at around mid-2014. The ongoing YFV outbreak in Brazil disseminated in the Southeast region following two independent routes that seems to be linked to the Paraná and Southeast Atlantic hydrographic basins, comprising densely populated regions. The spread of the YFV outside the Amazon and *Cerrado* biomes following primary hydrographic watersheds comprising large metropolis stresses the imperative importance of the continuous monitoring of YFV coupled with in depth phylogeographic analysis to aid decision-making Health authorities for effective prophylactic and control policies aiming the increase of vaccination coverage to avoid the YFV transmission in densely populated urban centers.

## 8 Author Contributions

Conceived the study: ED, GB, MCB, RLO

Carried out the collection of biological specimens: FVSA, MSASN Provided samples: EGK

Identification of mosquito species: IB, MSASN

Carried out viral RNA extraction from the biological specimens and the diagnosis by RT-PCR: AFB, FVSA, IB, LFFS, MGC, RMM

Inoculation of biological specimens in cell culture: LFFS, LMSR

Performed rapid viral RNA extraction and genome sequencing: AACS, IPR, MMG Analyzed the genome sequences: AACS, MCB, MMG, NDF

Performed phylogenetic/phylogeographic analysis: ED, GB

Prepared figures, tables and/or supplementary material: ED, GB, MCB, MMG, NDF, RLO Prepared the manuscript: ED, FVSA, GB, IPR, MCB, MMG, RLO, EGK, ACPV Gathered, systematized and illustrated epidemiological records: APR, DGR, FVSA

All authors critically read and approved the final version of the manuscript.

## 10 Acknowledgments

To Vinicius Lemes da Silva, Myriam A. F Campos, Angélica Bastos, Yulla Fernandes, Marcelo Santalucia (Secretaria de Saúde de Goiás), Romário Gabriel Aquino (Environmental Surveillance, Secretaria Municipal de Saúde de Angra dos Reis), Rodrigo F. C. Said (Secretria de Saúde de Minas Gerais), Cláudia Aarestrup, Livia Passarela, Adalberto Mitterofhe, Milton F. Castro, Adilson C. Lima (Secretarias Municipais de Saúde de Juiz de Fora e Belmiro Braga), Omar Figueiredo Neto (Environmental Surveillance, Secretaria Municipal de Saúde de Valença), Marilza L. Lange, Luciano L. Salles, Núcleo de Entomologia e Malacologia do Espírito Santo (NEMES), Tercius Barrada (Parque Estadual da Ilha Grande), Gilsa Aparecida P. Rodrigues (Secretaria de Saúde do Estado do Espírito Santo), Gilton Luiz Almada (Centro de Informação Estratégica de Vigilância em Saúde-ES), Mário Sérgio Ribeiro e Patrícia Menegueti (Secretaria de Saúde do Estado do Rio de Janeiro), Prefeitura de Macaé, Centro de Estudos Ambientais e Desenvolvimento Sustentável (CEADS/UERJ), Marcelo Celestino dos Santos, Mauro M. Muniz, Marcelo Q. Gomes, Teresa F. Silva-do-Nascimento for the help in obtaining the viral samples and support in the field work.

## 12 Data Availability Statement

All new YFV genomes generated for this study were submitted to GenBank and their accession numbers are included in the manuscript. The information of all YFV genomes used in this study is provided in the Supplementary Table 1.

## Supplementary information

**Supplementary Table 1.**
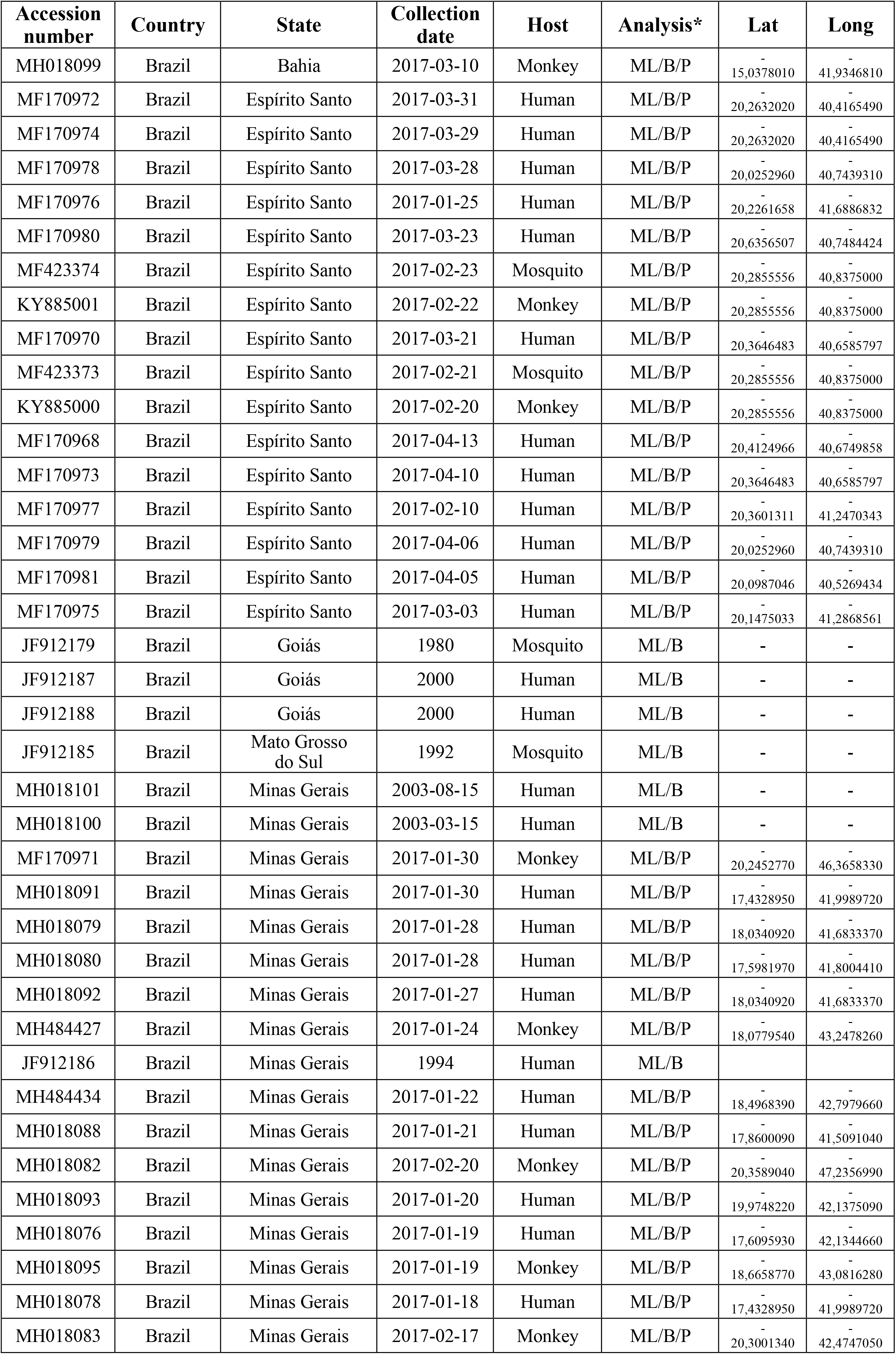
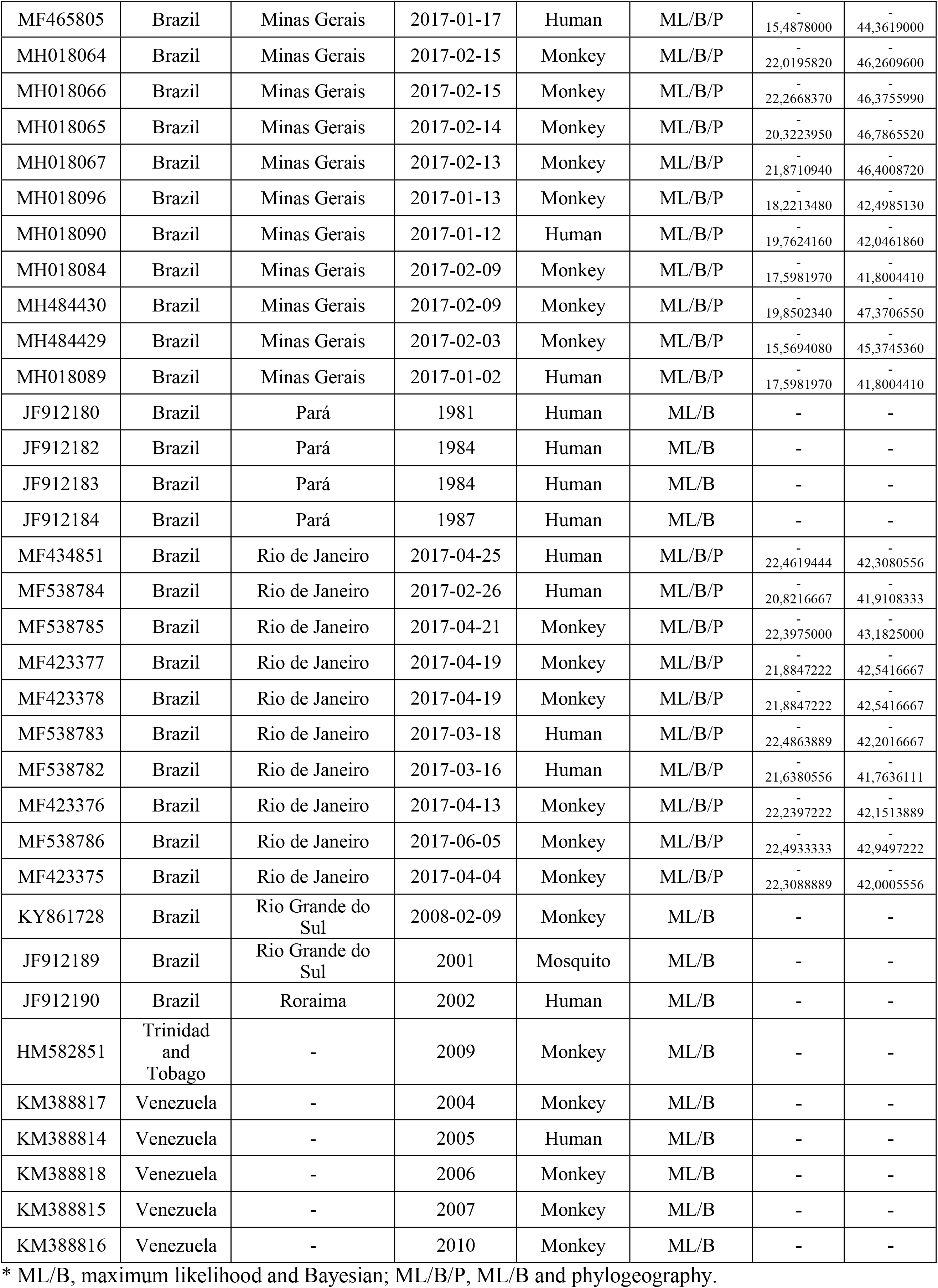
Information about all YFV complete genomes used in this study. lineage using different combinations of clock rate priors and phylogeographic models

**Supplementary Table 2.**
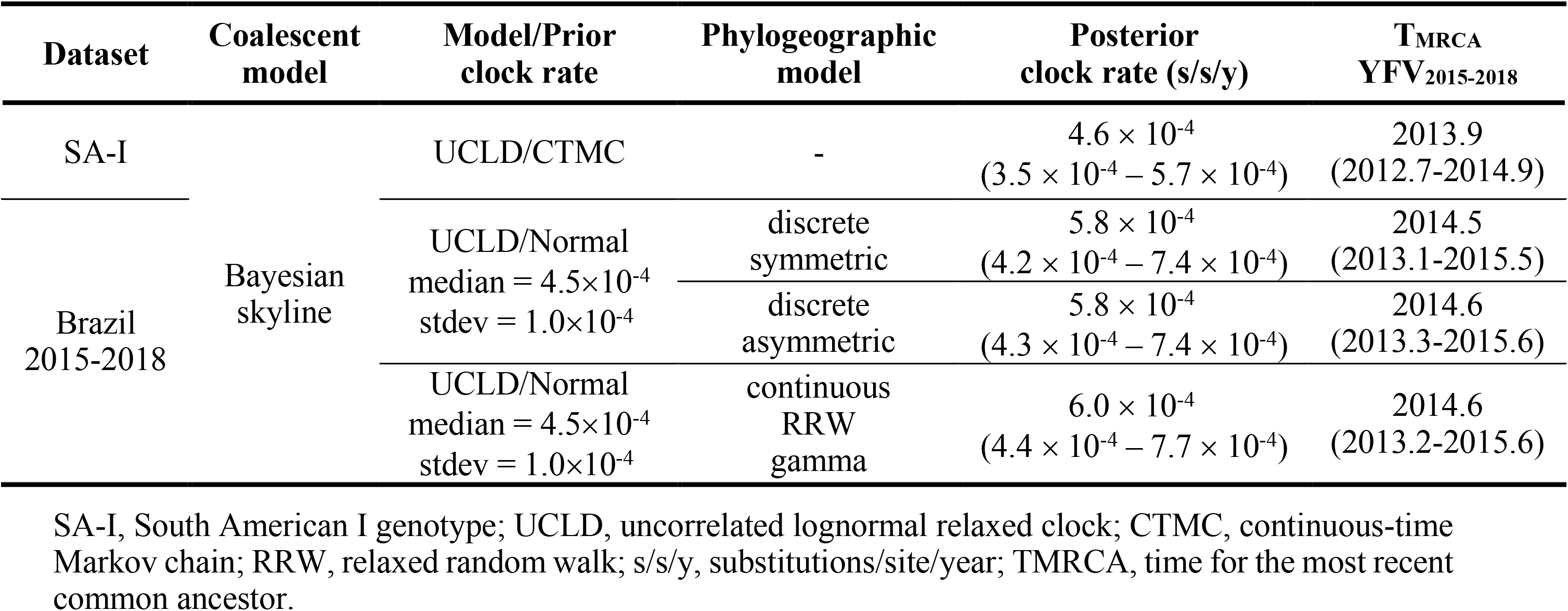
Molecular cock rates and T_MRCA_ obtained for the YFV_2015-2018_ lineage using different combinations of clock rate priors and phylogeographic models

**Supplementary Table 3.** Amino acid molecular signature present in 2016-2018 YFV samples in Southeastern Brazil.

| 2016-2018 YFV genomes<br>(GenBank accession number) | Polyprotein position |  |  |  |  |  |  |  |  |
| --- | --- | --- | --- | --- | --- | --- | --- | --- | --- |
|  | 108 | 1572 | 1605 | 2607 | 2644 | 2679 | 2803 | 3149 | 3215 |
| KY885000; KY885001; MF170968;<br>MF170970; MF170971; MF170972;<br>MF170973; MF170974; MF170975;<br>MF170976; MF170977; MF170978;<br>MF170979; MF170980; MF170981;<br>MF423373; MF423374; MF423375;<br>MF423376; MF423377; MF423378;<br>MF434851; MF465805; MF538782;<br>MF538783; MF538784; MF538785;<br>MF538786; MH018065; MH018066;<br>MH018067; MH018076; MH018078;<br>MH018079; MH018080; MH018082;<br>MH018083; MH018090; MH018091;<br>MH018092; MH018095; MH018096;<br>MH018099; MH484427; MH484429;<br>MH484430; MH484434; MK333798;<br>MK333799; MK333800; MK333801;<br>MK333802; MK333805; MK333806;<br>MK333807; MK333808; MK333809 | I | D | K | R | I | S | S | A | S |
| MH018088; MH018084; MH018089;<br>MH018064 | I | D | K | Q | I | S | S | A | S |
| MH018093 | I | E | K | Q | I | S | S | A | S |

**Supplementary Table 4.** Comparison of discrete spatial models fit to the 2015-2018 Brazilian YFV dataset.

| **Model** | **PS**<br>**Log ML** | **Models**<br>**compared** | **Log BF** | **SS**<br>**Log ML** | **Models**<br>**compared** | **Log BF** |
| --- | --- | --- | --- | --- | --- | --- |
| Sym | -16226.06 | - | - | -16226.54 | - | - |
| Asym | -16225.78 | Asym/Sym | 0.28 | -16226.60 | Sym/Asym | 0.06 |

Sym, symmetric; Asym, asymmetric, PS, path sampling, SS, stepping-stone; ML, marginal likelihood; BF, Bayes factor.

**Supplementary Table 5.** Comparison of continuous spatial models fit to the 2015-2018 Brazilian YFV dataset and estimate of YFV_2015-2018_ lineage dispersal rate under the different models.

|  | **Homogeneous**<br>**Brownian diffusion** | **Heterogeneous**<br>**RRW Cauchy** | **Heterogeneous**<br>**RRW Gamma** | **Heterogeneous**<br>**RRW Lognormal** |
| --- | --- | --- | --- | --- |
| PS - Log ML<sup>a</sup> | -16339 | -16321 | -15777 | -16324 |
| PS – Log BF<sup>b</sup> | 562 | 544 | Fittest model | 547 |
| SS - Log ML<sup>a</sup> | -16340 | -16321 | -14807 | -16324 |
| SS – Log BF<sup>b</sup> | 1533 | 1514 | Fittest model | 1517 |
| Dispersal rate<br>(km/day)<sup>c</sup> | 0.7<br>(0.4 – 1.0) | 0.7<br>(0.5 - 1.0) | 0.8<br>(0.5 - 1.0) | 0.7<br>(0.5 - 1.0) |

<sup>a</sup> Log marginal likelihood (ML) estimates for the different continuous phylogeographic models obtained using the path sampling (PS) and stepping-stone sampling (SS) methods. <sup>b</sup> The Log Bayes factor (BF) is the difference of the Log ML between of alternative (H1) and null (H0) models (H1/H0). Log BF<sub>s</sub> > 3 indicates that model H1 is more strongly supported by the data than model H0. <sup>c</sup> Posterior mean and 95% HPD (in parenthesis) estimates of the dispersal rate.

**Supplementary figure 1.**
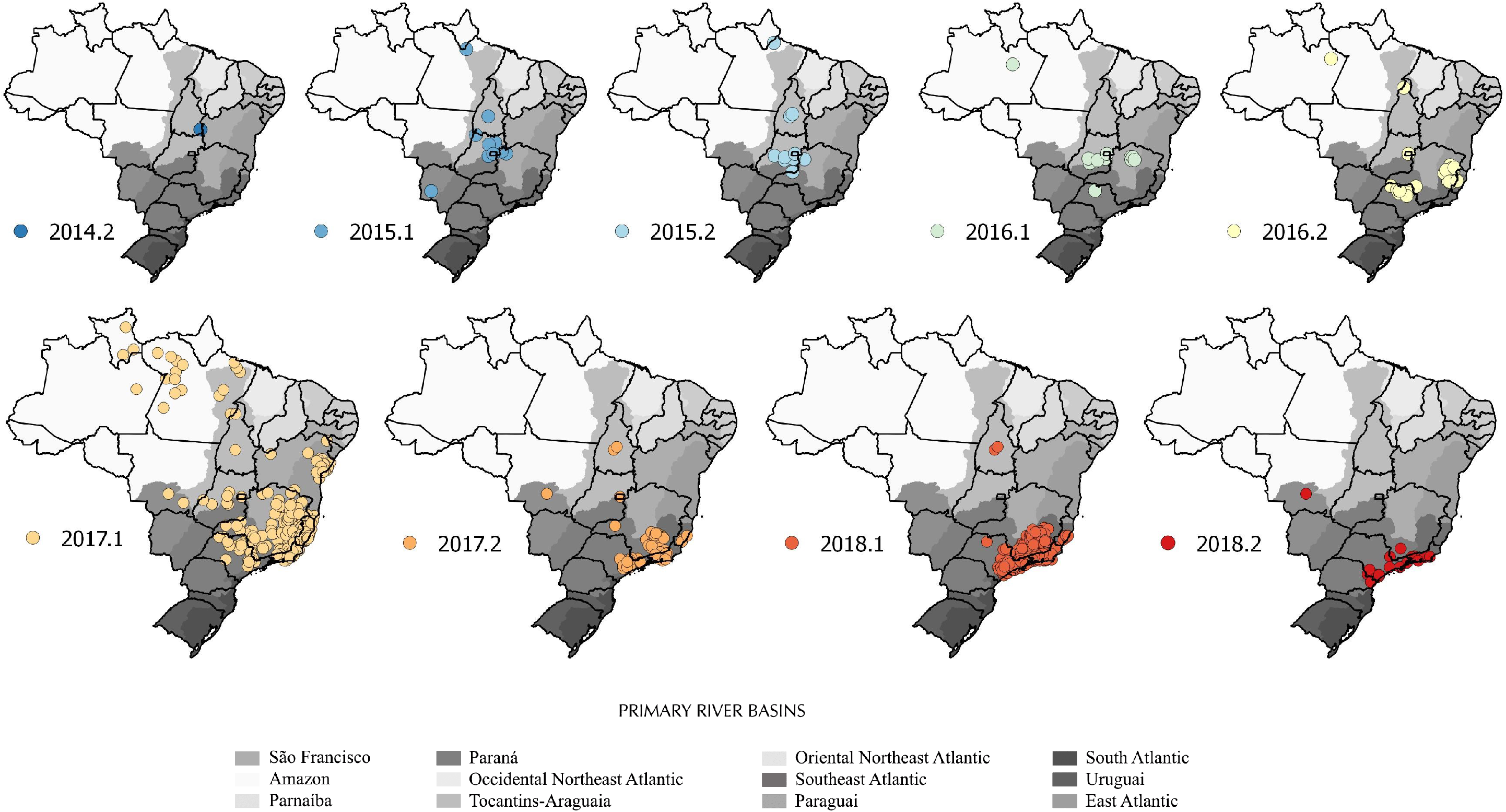
Half-yearly based spatial spread of YFV in Brazil from the 2^nd^ semester 2014 (2014.2) to 2^nd^ semester 2018 (2018.2) according to epidemiological records. Dots corresponds to the centroid of municipalities where human and/or non-human YFV infections were laboratory confirmed. The gray-shaded background underlines primary Brazilian river basins. Source: SVS-Brazilian Ministry of Health.

**Supplementary Figure 2.**
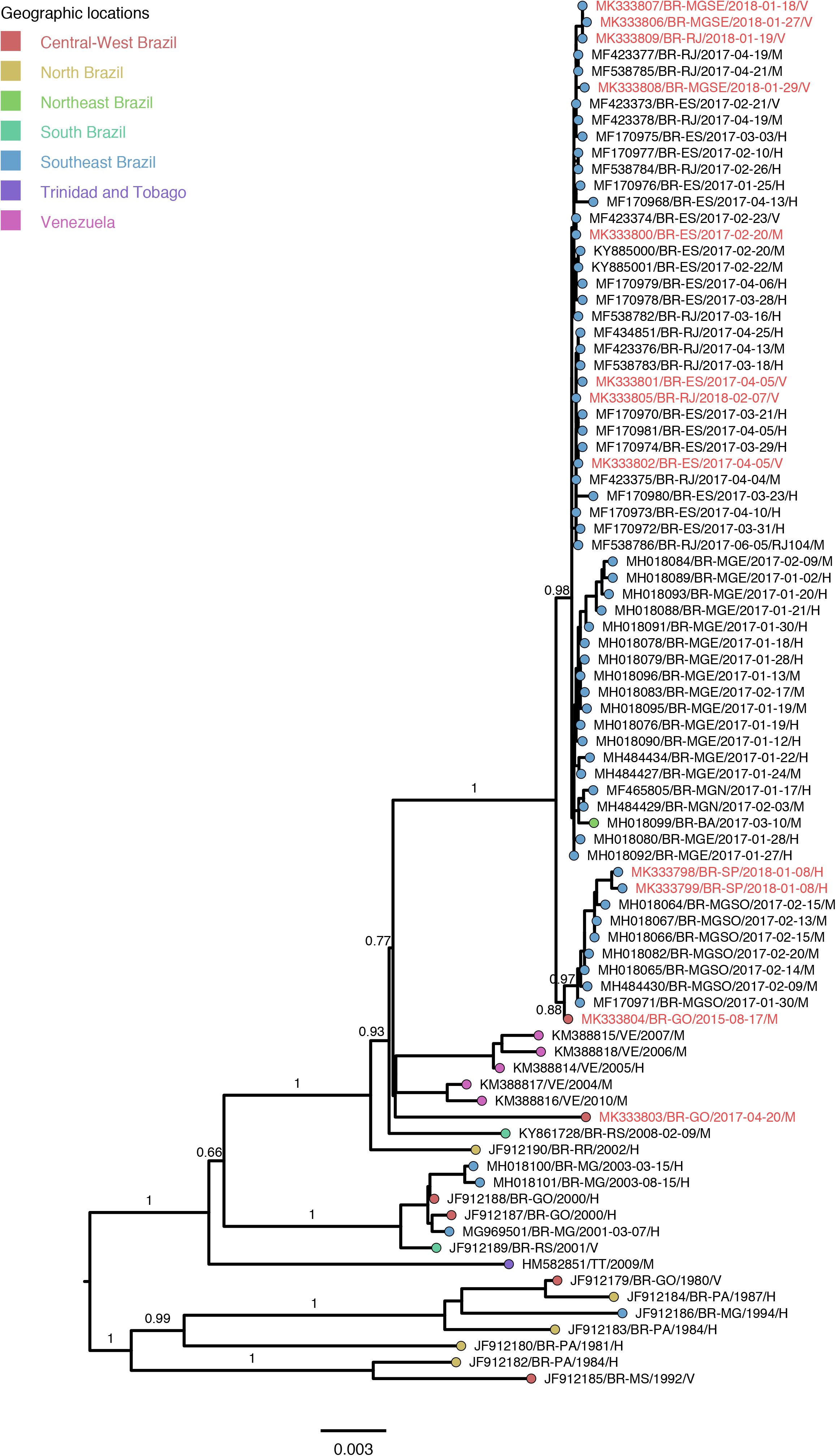
Maximum likelihood phylogeny of YFV South American I complete genome sequences. The aLRT support ue of key nodes are indicated. Tip circcles are colored following the legend at top left indicating the Brazilian region or country ampling. The branch lengths are drawn to scale with bar at the bottom indicating nucleotide substitutions per site. Tips names of the sequences from this study were colored red. The last letter of the sequence name indicate the host: H - human; mosquito vector; M - non-human primate.

**Supplementary Figure 3.**
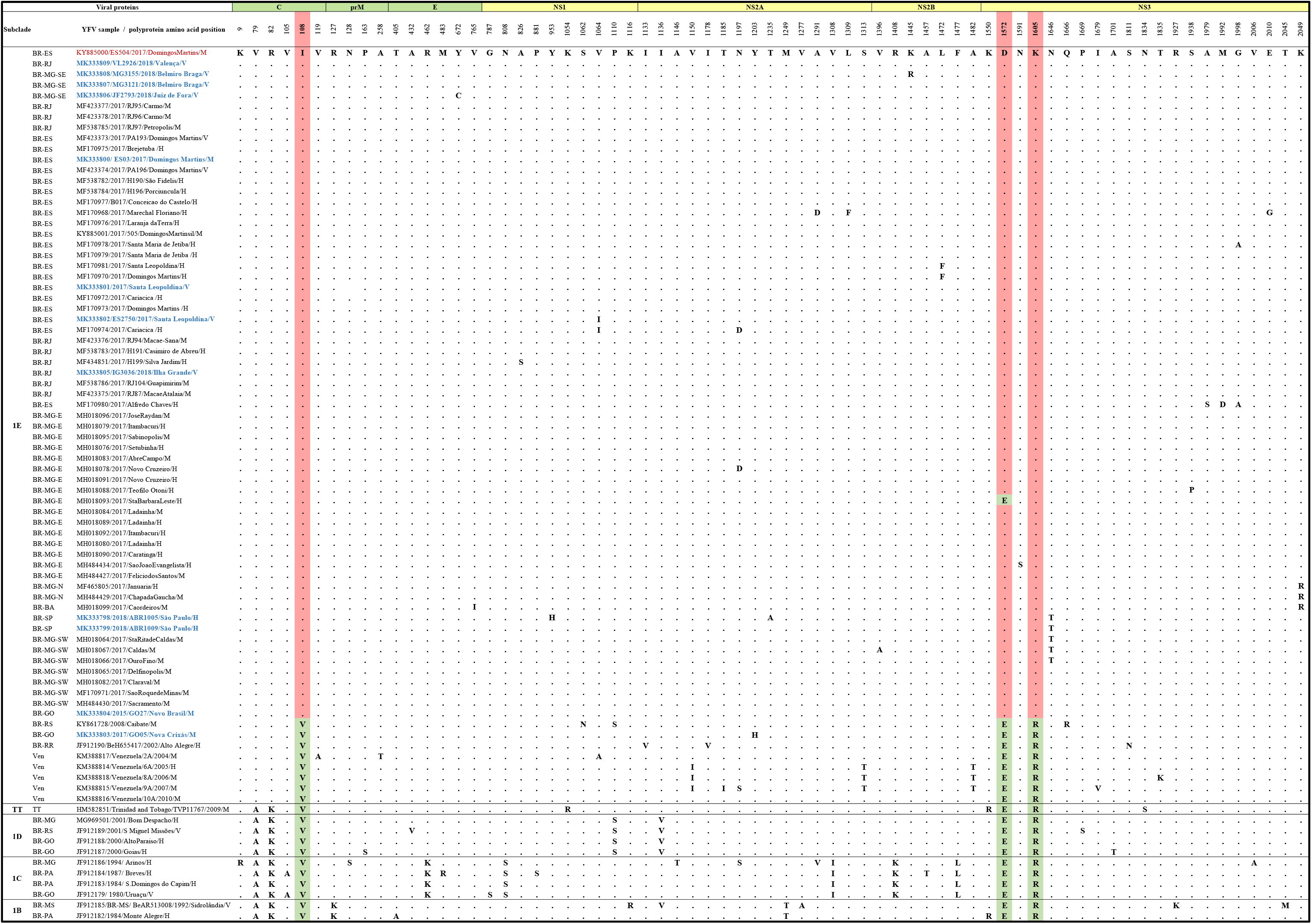
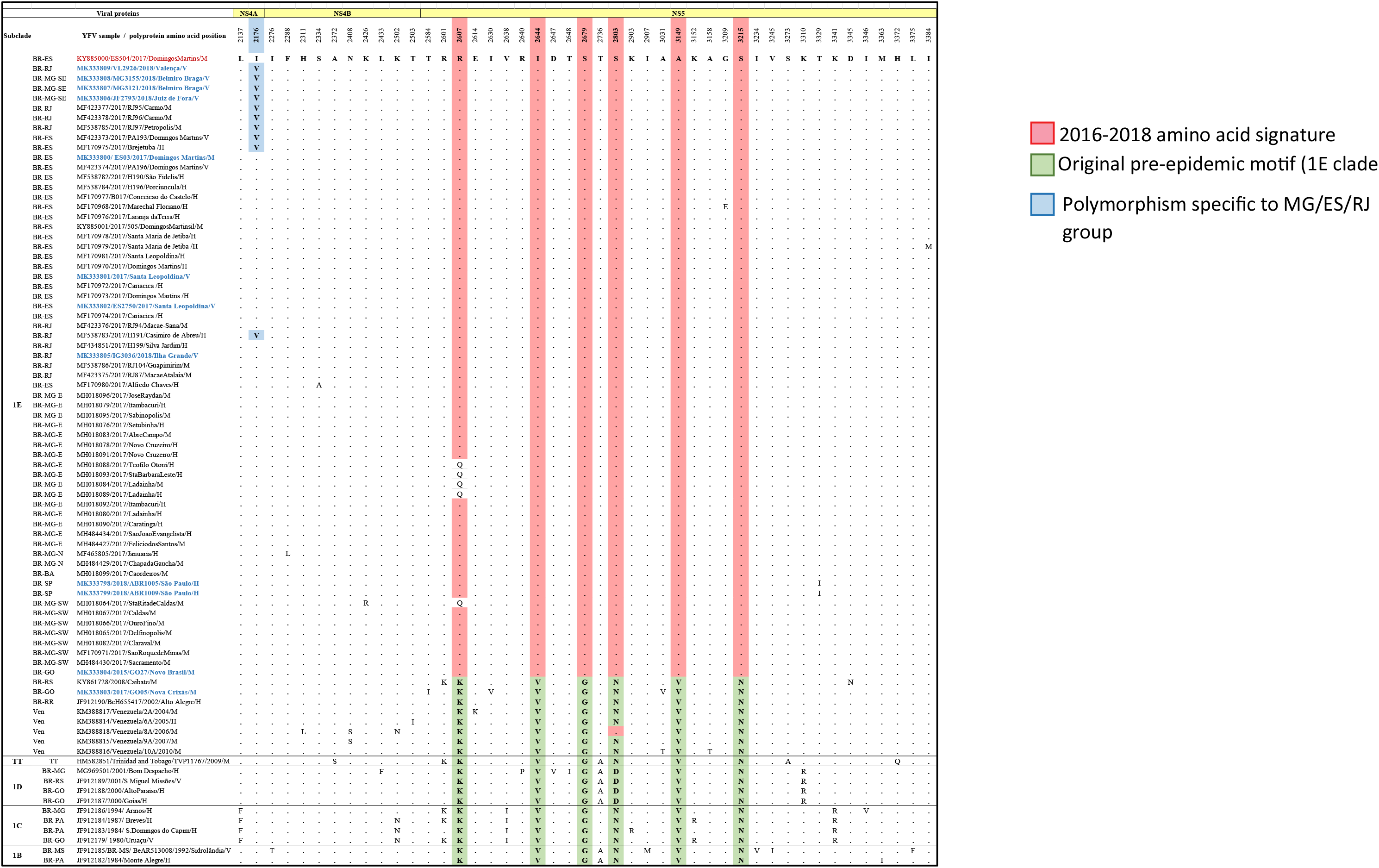
Amino acid polymorphisms in YFV in the precursor polyprotein

